# Sequences enriched in Alu repeats drive nuclear localization of long RNAs in human cells

**DOI:** 10.1101/189746

**Authors:** Yoav Lubelsky, Igor Ulitsky

## Abstract

Long noncoding RNAs (lncRNAs) are emerging as key players in multiple cellular pathways, but their modes of action, and how those are dictated by sequence remain elusive. While lncRNAs share most molecular properties with mRNAs, they are more likely to be enriched in the nucleus, a feature that is likely to be crucial for function of many lncRNAs, but whose molecular underpinnings remain largely unclear. In order identify elements that can force nuclear localization we screened libraries of short fragments tiled across nuclear RNAs, which were cloned into the untranslated regions of an efficiently exported mRNA. The screen identified a short sequence derived from Alu elements and found in many mRNAs and lncRNAs that increases nuclear accumulation and reduces overall expression levels. Measurements of the contribution of individual bases and short motifs to the element functionality identified a combination of RCCTCCC motifs that are bound by the abundant nuclear protein HNRNPK. Increased HNRNPK binding and C-rich motifs are predictive of substantial nuclear enrichment in both lncRNAs and mRNAs, and this mechanism is conserved across species. Our results thus detail a novel pathway for regulation of RNA accumulation and subcellular localization that has been co-opted to regulate the fate of transcripts that integrated Alu elements.

## Introduction

Long noncoding RNAs are increasingly recognized as important players in various biological systems^1^, but their modes of action, and how those are encoded in their linear sequences, remain largely unknown. In most known aspects of their molecular biology, lncRNAs closely resemble mRNAs; both transcript types start with a 5' cap, end with a poly(A) tail, and are typically spliced. As a group, lncRNAs tend to be enriched in the nuclear fraction, whereas most mRNAs are overtly cytoplasmic^2^, although several studies have found that hundreds of mRNAs in various cell types are retained in the nucleus, potentially contributing to robustness of protein expression levels in spite of bursty transcription^3,4^. It is thus conceivable that some mechanisms that promote nuclear enrichment are shared between lncRNAs and some mRNA transcripts.

As there are no well-characterized pathways for the import of long RNAs into the nucleus, nuclear enrichment likely corresponds to either inhibition of export, leading to nuclear retention, or to destabilization of transcripts in the cytoplasm. However, the detailed mechanisms and sequence elements that can direct nuclear enrichment remain unknown for the majority of long RNAs, with a few exceptions^5–8^. Here, we use a high-throughput screen to identify a sequence element present in both lncRNA and mRNA transcripts, which is associated with widespread nuclear enrichment. We also identify the protein that binds this element, HNRNPK, and show that it is required for the nuclear enrichment of the bound RNAs, potentially in cooperation with other factors or pathways.

## Results

We hypothesized that the nuclear localization observed for some of the lncRNAs and mRNAs is encoded in short sequence elements capable of autonomously dictating nuclear enrichment in an otherwise efficiently exported transcript. In order to systematically identify such regions, we designed a library of 5,511 sequences of 109 nt, composed of fragments that tile the exonic sequences of 37 human lncRNAs, 13 3' UTRs of mRNAs enriched in the nucleus in mouse liver^3^, and four homologs of the abundant nuclear lncRNA MALAT1^9^ (**Tables S1 and S2**). These tiles were cloned into the 5' and 3' UTRs of AcGFP mRNA (NucLibA library, **Figure 1A**). Offsets between consecutive tiles were typically 25 nt (10 nt in MALAT1 and TERC and 50 nt in the longer XIST and MEG3 genes). After transfection into MCF7 cells in triplicates, we sequenced the inserts from GFP mRNAs from whole cell extract (WCE), nuclear (Nuc), and cytoplasmic (Cyt) fractions, as well as from the input plasmids (>10 million reads per sample, **Table S3**). Normalized counts of unique molecular identifiers (UMIs, **Table S2**) were used to evaluate the effect of each inserted tile on expression levels and subcellular localization of the GFP mRNA (**Table S4**).

**Figure 1.**
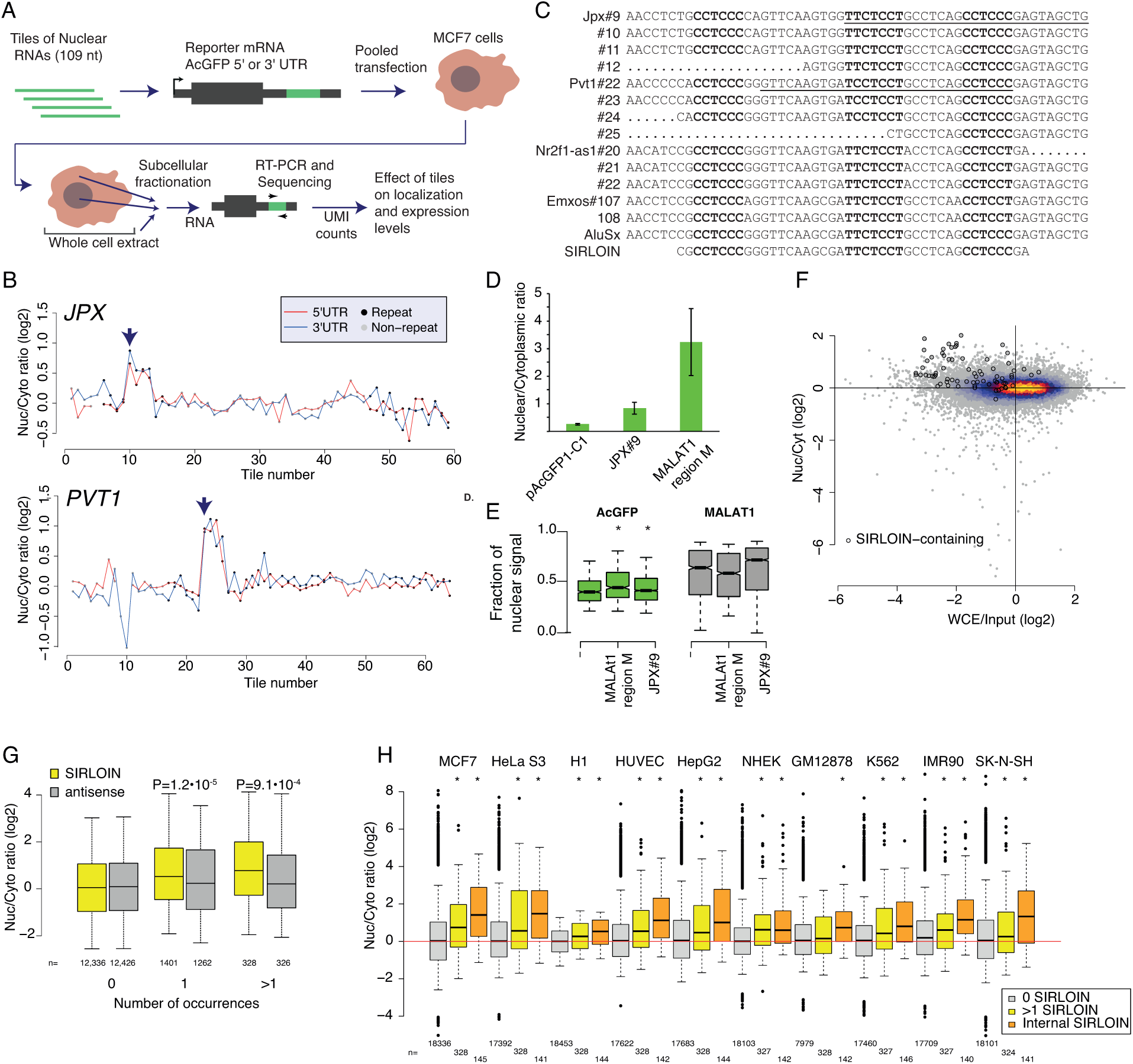
**(A)** Outline of the experimental approach. **(B)** Effects of all the tiles in JPX and PVT1 lncRNAs on nuclear/cytoplasmic expression ratios when cloned into the indicated region of AcGFP. Tiles overlapping repetitive elements are in black and other tiles are in gray. **(C)** Sequence alignment of 13 of the most effective tiles, the consensus sequence of AluSx repeat family, and the SIRLOIN element. C/T-rich hexamers are in black, and the regions in JPX#9 and PVT1#22 that were mutagenized in NucLibB are underlined. **(D)** qRT-PCR determination of the abundance of AcGFP mRNA with the indicated fragments cloned into the 3’ UTR; nuclear/cytoplasmic ratios are shown. **(E)** Imaging flow cytometry of AcGFP and MALAT1 mRNA, the fraction of the signal overlapping with DAPI signal out of the total signal intensity is displayed. Asterisks denote P<0.02 (Wilcoxon test). **(F)** Correlation between the effects of individual tiles on expression levels (x-axis, WCE is whole cell extract) and nuclear/cytoplasmic ratios (y-axis). SIRLOIN-containing elements are in circles. **(G)** Nuclear/cytoplasmic expression ratios for RNAs containing the indicated number of SIRLOIN elements or SIRLOIN reverse complement (antisense) in MCF7 cells ENCODE data. P-values are for comparisons between the RNAs with the indicated number of SIRLOINs in the sense vs. antisense orientation. **(H)** Nuclear/cytoplasmic ratios in each of ten ENCODE cell lines, comparing transcripts without SIRLOIN elements to those with at least two elements and to those with at least one element in an internal exon. Asterisks indicate P<0.01 (Wilcoxon test) when comparing the indicated group to transcripts without SIRLOIN elements.

In order to identify high-confidence effects, we focused on consecutive tiles with a consistent effect on localization. We identified 19 regions from 14 genes spanning 2–4 overlapping tiles, with each of the tiles associated with a >30% nuclear enrichment (**Table S5**). The three regions where the tiles had the highest enrichments originated from the lncRNAs JPX, PVT1, and NR2F1-AS1, and those had similar activity when placed in either the 3' or 5' UTR of the GFP mRNA (**Figure 1B and Figure S1**). These tiles overlapped Alu repeat sequences inserted in an ‘antisense’ orientation, and the overlap region between the active patches converged on a 42 nt fragment that contained three stretches of at least six pyrimidines (C/T), two of which were similar to each other and matched the consensus RCCTCCC (R=A/G). We named this 42 nt sequence element SIRLOIN (SINE-derived nuclear RNA LOcalizatIoN) (**Figure 1C**). We validated specific SIRLOIN-containing tiles cloned individually for their ability to drive nuclear enrichment of the GFP mRNA using qRT-PCR (**Figure 1D**, the MALAT1 ~600 nt “region M”^5^ is used as positive control) and imaging flow cytometry with the PrimeFlow^™^ RNA assay (**Figure 1E**). JPX#9 was also validated to drive nuclear localization by another group using single molecule FISH^10^ (John Rinn, personal communication).

Interestingly, despite the fact that overall, there was no significant correlation between the effects of individual tiles on expression levels and on localization (Spearman R=0.001, P=0.82), SIRLOIN-containing tiles were associated with consistently lower AcGFP RNA expression levels (Spearman R=-0.31 between effects on localization and expression, P=8.5·10^-3^, **Figure 1F**). SIRLOIN elements thus affect both the localization and the expression levels of transcripts.

In order to test whether SIRLOIN elements were globally associated with nuclear localization in long RNAs, we analyzed RNA-seq data from human cell lines profiled by the ENCODE project^2^. SIRLOIN elements were associated with nuclear enrichment in MCF7 cells (**Figure 1G**), and two or more SIRLOIN elements were associated with a significant (P<0.01, Wilcoxon test) nuclear enrichment in nine out of ten ENCODE cell lines (**Figure 1H**). Interestingly, SIRLOIN elements in internal exons (as in the endogenous context of the SIRLOINs in JPX, PVT1, and NR2F1-AS1) were associated with stronger nuclear enrichment, and in all ten cell lines, a single SIRLOIN element in an internal exon was associated with a significant nuclear enrichment (**Figure 1H**), with similar trends observed for both mRNAs and lncRNAs (**Figure S2**).

To better characterize the effect of SIRLOIN on RNA localization in human lncRNAs and mRNAs, we designed and cloned a second library, NucLibB (**Tables S6-8**), that included a large number of sequence variations within two 30 nt fragments of JPX#9 and PVT1#22 that were two of the most highly effective tiles in NucLibA. The variants included mutations of single bases or consecutive sequence stretches, repetitions of segments of various lengths, and shuffled controls, all within the same fixed insert length of 109 nt. We also included tiles from six lncRNAs and five human mRNAs that overlapped SIRLOIN elements. These included four mRNAs with inverted pairs of Alu elements (“IRAlu”), that were previously shown to drive nuclear retention with a possible connection to A-to-I editing^11^, though the generality of this phenomenon has been questioned^12^. Since in NucLibA we observed that tiles cloned into either the 5' or 3' UTRs exerted similar effects, we cloned NucLibB only into the 3' UTRs of the AcGFP mRNA.

Using NucLibB we validated the nuclear-enrichment activity of the JPX and PVT1 SIRLOINs, and identified 18 additional SIRLOIN-overlapping regions from five lncRNAs and five mRNAs that met the same criteria used in NucLibA (at least two consecutive tiles causing nuclear enrichment of >30%, **Table S9**). In some cases, several regions from the same transcript were effective in mediating nuclear localization (**Figure 2A and S3**) and SIRLOIN-matching elements were consistently active (**Figure 2B**). The new SIRLOIN-containing sequences identified in NucLibB also had a substantial negative impact on expression levels, and effects on localization strongly correlated with effects on expression levels (Spearman R=–0.6, P<10^-16^, **Figure S4**). Interestingly, among the SIRLOIN-containing sequences in NucLibB, those that had a SIRLOIN closer to the 3' end were more enriched in the nucleus (R=0.34, P=0.037) and more poorly expressed (R=-0.55, P=0.0004), suggesting that SIRLOIN context affects its activity (**Figure 2C**).

**Figure 2.**
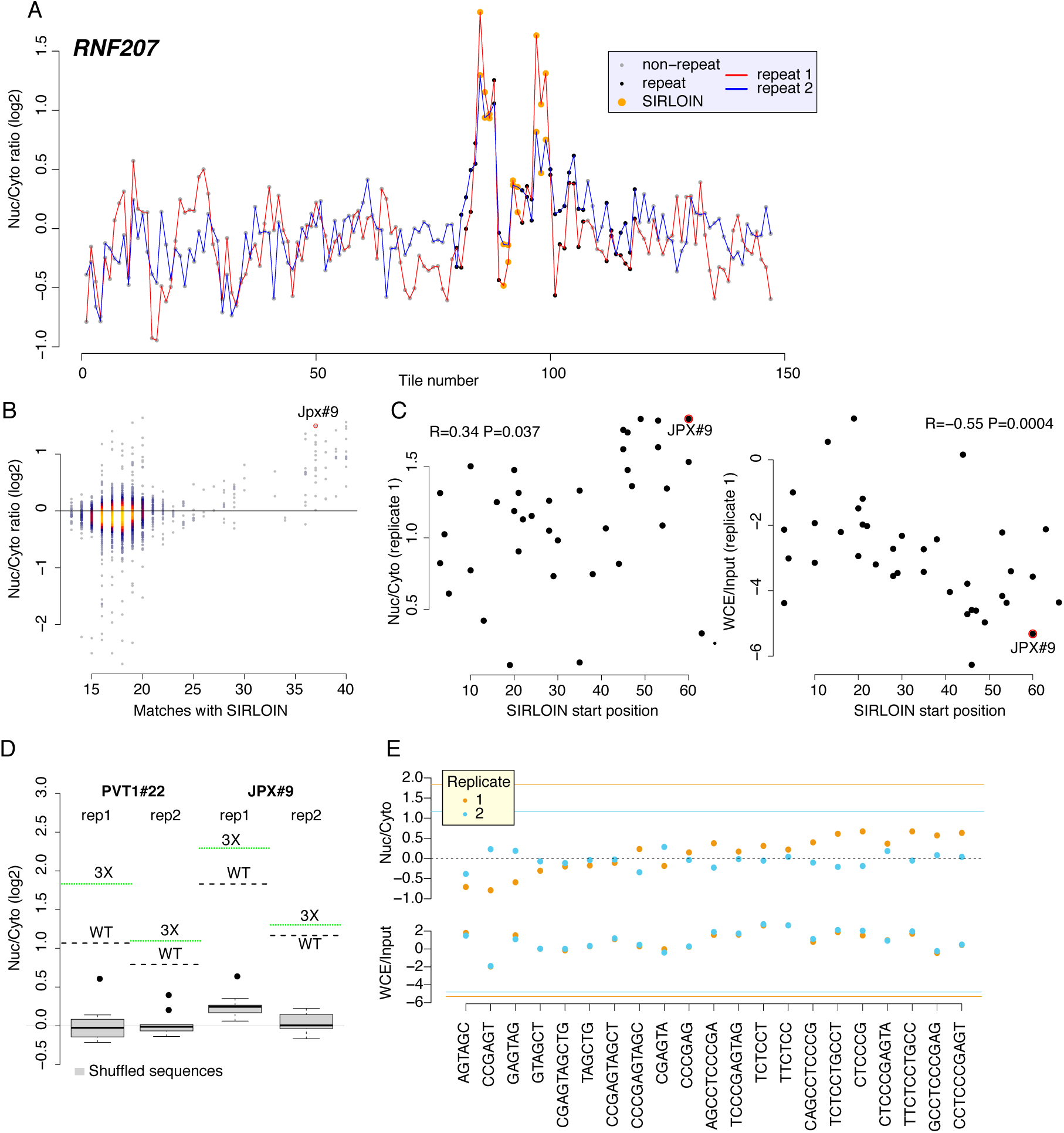
**(A)** Effects of the tiles in RNF207 mRNA on nuclear/cytoplasmic expression ratios when cloned into the 3’UTR of AcGFP. Tiles overlapping repetitive elements are in black, tiles overlapping the SIRLOIN element are in orange, and other tiles are in gray. **(B)** Correlation between similarity to a SIRLOIN element (number of perfectly matching bases, without allowing indels) and Nuc/Cyto ratios in NucLibB. **(C)** Correlation between the position of the SIRLOIN element within the tile and the effect of the tile on localization (left) and expression levels (right). Spearman’s correlation coefficients and p-values are indicated. **(D)** Box plots show effects on Nuc/Cyto ratio of ten shuffled, dinucleotide-preserving sequences of the indicated tiles in each of two replicates. Horizontal lines show the effects of the WT sequence of the tile as well as sequences containing 3 core parts of the SIRLOIN element (underlined in Figure 1C). **(E)** Effects of 109 nt sequences containing repeats of the indicated 6‐ or 10-mers from JPX#9 tile separated by AT dinucleotides on localization (top) and expression levels (bottom). Horizontal lines show the effects of the WT JPX#9 sequence.

NucLibB included shuffled sequences of JPX#9 and PVT1#22 (preserving their dinucleotide composition), and as expected, those did not impose nuclear enrichment (**Figure 2D**). Within the limits of 109 nt fragments in our library, we also tested the activity of sequences containing multiple SIRLOIN elements. Three repeats of core SIRLOIN parts from JPX#9 and PVT1#22 cloned into one tile caused substantially stronger nuclear enrichment than sequences containing only one SIRLOIN element (**Figure 2D**). NucLibB also contained sequences composed of repetitions of individual 6‐ or 10-mers from the cores of JPX#9 and PVT1#22 SIRLOIN elements, separated by AT dinucleotides. While we observed more nuclear-enrichment activity from the C/T-rich elements (**Figures 2E and S5**), such tiles exerted very limited effects on either nuclear enrichment or expression levels, suggesting that secondary structure or sequences beyond the C/T-rich motifs are important for SIRLOIN function.

We next analyzed the effects of sequence changes on SIRLOIN activity. Single point mutations to purines (A/G) in the second RCCTCCC motif of JPX#9 were sufficient to abolish the effect of the sequence on both localization and expression levels, whereas C→T mutations in that region had little effect (**Figure 3A and Figure S6A**). More extensive changes, alternating A↔T and G↔C, were deleterious also when made outside of the RCCTCCC motif, suggesting that additional parts of SIRLOIN are essential for its function, but can tolerate single base changes (**Figure 3B**). The PVT1#22 sequence was interestingly more resilient to changes, and single point mutations had a limited effect on activity (**Figure S6B**). More extensive changes to PVT1#22, including mutating 4 bases in the 3' part of its SIRLOIN element, were sufficient to abolish activity (**Figure S6C**). We conclude that the second RCCTCCC motif is the most important part of the SIRLOIN element, but that other SIRLOIN regions also contribute to its function.

**Figure 3:**
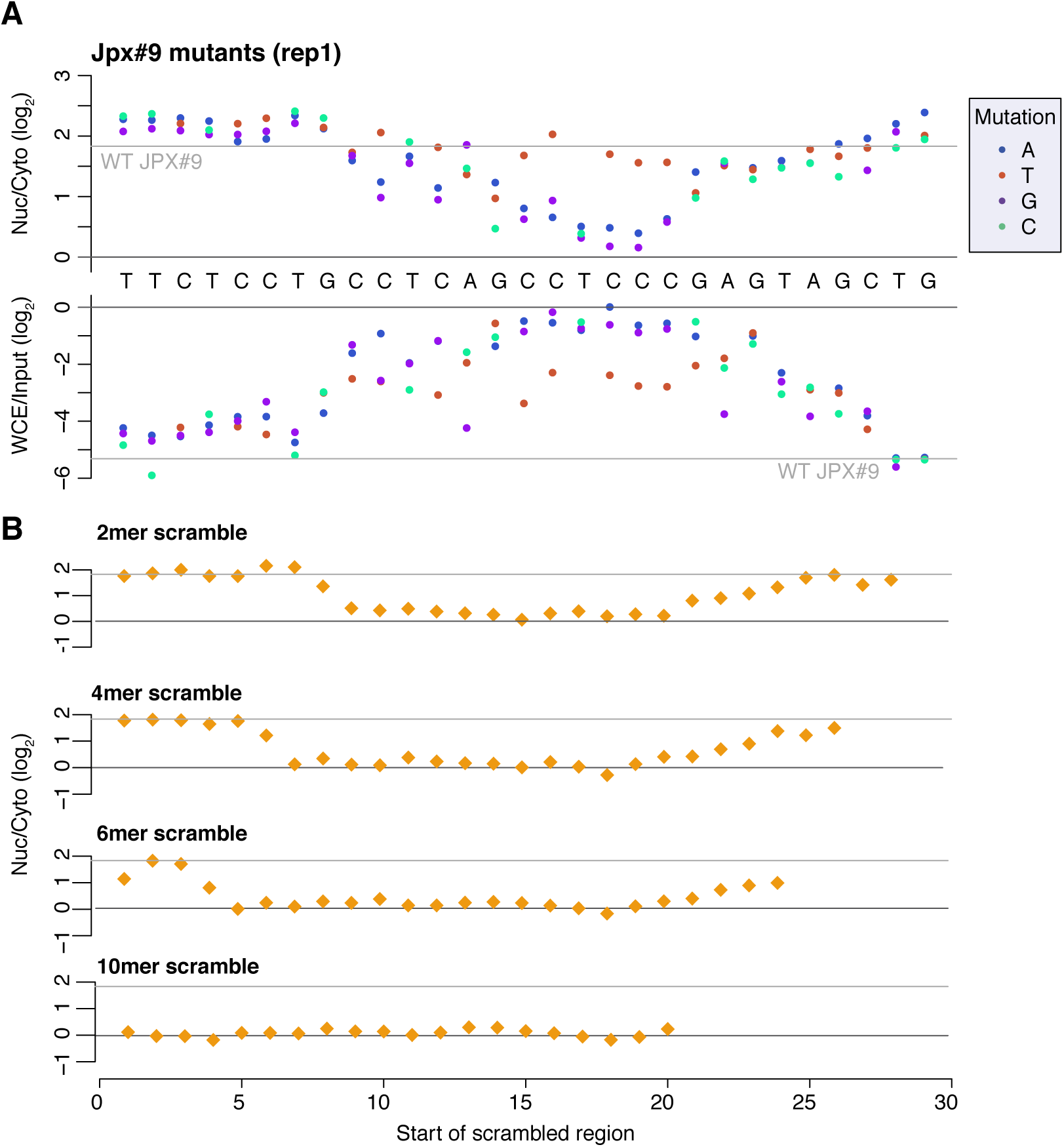
**(A)** Nuclear/cytoplasmic ratios (top) and expression levels (bottom) for sequence variants that are all identical to JPX#9 except for the indicated base change. Horizontal gray line indicates the corresponding level for the wild type JPX#9 sequence. WT sequence is shown between the two plots. **(B)** Nuclear cytoplasmic ratios for sequences where the indicated number of bases were scrambled with C↔G and A↔T changes. The position of the point indicates the first base of the region that was scrambled.

RNA editing was suggested to play a causal role in nuclear retention of IRAlu-containing RNAs^7^. However, the RNAs we studied are not predicted to form long intramolecular double stranded RNA (dsRNA) regions that would be effective substrates for A-to-I editing (for example, see **Figure S7** for the predicted secondary structures of AcGFP mRNAs with JPX#9 and PVT1#22 in their 3' UTRs). In order to evaluate the extent of RNA editing in our library inserts, we analyzed the signatures of A-to-I RNA editing in the sequencing data of NucLibB. A→G mutations indicative of A-to-I editing were more prevalent than other mutations in cDNA reads when normalized to the input plasmids (**Figure S8A**).

The overall editing levels of SIRLOIN-containing sequences were typically low (<3%), but higher than the levels in other NucLibB sequences (**Figure S8B-D**), and fragments with higher editing levels were enriched in the nucleus and expressed at significantly lower levels (**Figure S8E**, P=1.8·10^-4^ and P=3.5·10^-9^ for localization and expression, respectively). However, it is unlikely that A-to-I editing plays a causal role in nuclear enrichment of SIRLOIN-containing sequences, as the per-position editing levels were very low, and >97% of reads sequenced from the nuclear fractions and mapping to SIRLOIN-containing tiles did not contain any edited bases. It is possible that the plausibly longer duration that transcripts enriched in the nucleus spend in the nucleus (compared to those that are efficiently exported) lead to a higher probability of RNA editing, which in the case of SIRLOIN-containing tiles may act on inter-molecular dsRNA formed with RNAs encompassing ‘sense strand’ Alu elements; alternatively, some other structure or element associated with SIRLOIN elements affects their susceptibility to undergo editing.

We next hypothesized that SIRLOIN elements act through interactions with specific RNA-binding proteins. To this end, we queried the ENCODE eCLIP datasets for protein-RNA interaction sites specifically enriched in SIRLOINs (see Methods). eCLIP experiments for HNRNPK, an abundant nuclear RNA binding protein with known roles in the biology of specific lncRNAs, such as XIST and TP53COR1 (linc-p21)^13,14^, ranked first in this analysis (**Figure 4A**). Reassuringly, the motif enriched in the HNRNPK binding peaks throughout the transcriptome, as identified by GraphProt^15^, was a pyrimidine-rich sequence with a CCTCC core (**Figure 4B**), consistent with the known preferences of HNRNPK for C-rich sequences^16^, and matching the RCCTCCC sequence we identified as required for SIRLOIN function by the mutagenesis analysis. Moreover, the three KH RNA-binding domains of HNRNPK were previously shown to act cooperatively in binding sequences with triplets of C/T-rich regions^17^, fitting the sequence architecture of the SIRLOIN elements. GraphProt analysis also suggested the C/T-rich motif is preferentially bound in a structured context (**Figure 4B**), consistent with our observation that simple repeats of short motifs from SIRLOIN were not functional. Using RNA immunoprecipitation (RIP) with an HNRNPK-specific antibody, we validated that HNRNPK binds to AcGFP mRNA supplemented with SIRLOIN elements, but not to single-nucleotide mutants that were not effective in nuclear enrichment (**Figure 4C**).

**Figure 4.**
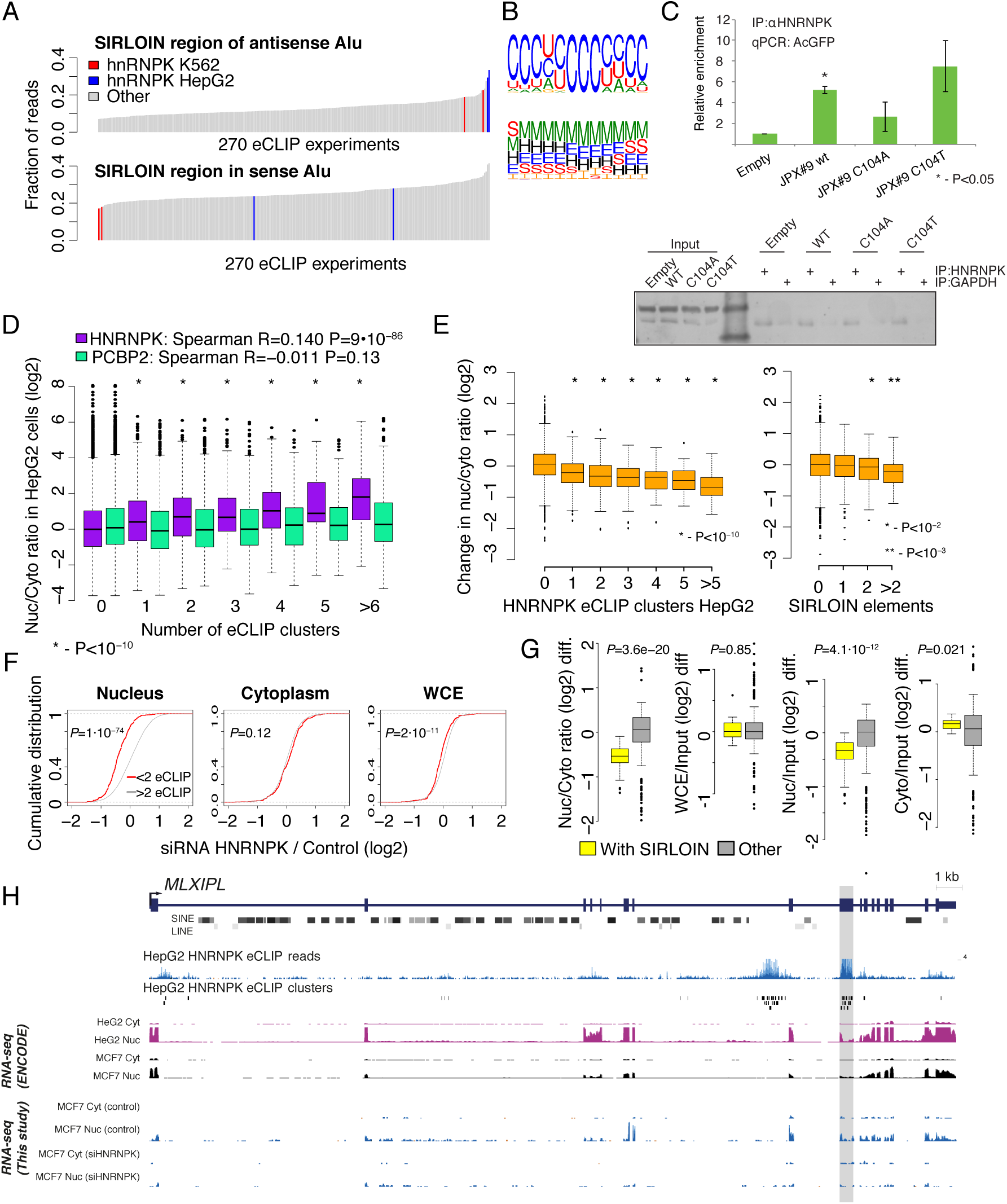
HNRNPK drives nuclear localization of SIRLOIN-containing transcripts. **(A)** Enrichment of HNRNPK in binding to the SIRLOIN element within Alu repeats (see Methods). **(B)** Sequence (top) and structure (bottom) logo of the motif enriched in HNRNPK eCLIP clusters (HepG2 cells, first replicate), as identified by GraphProt. Structural annotation: stems (S), external regions (E), hairpins (H), internal loops (I), multiloops (M) and bulges (B). **(C)** Enrichments of the AcGFP RNA with the indicated inserts following immunoprecipitation (IP) using an HNRNPK antibody, normalized to the GAPDH antibody. **(D)** Ratios of expression levels in the nucleus/cytoplasm for transcripts with the indicated number of eCLIP clusters for HNRNPK and PCBP2 in HepG2 cells. Asterisks indicate significant difference between the genes with the indicated number of clusters and genes without eCLIP clusters (Wilcoxon test). **(E)** Change in the nuclear / cytoplasmic ratio following HNRNPK knockdown for transcripts with the indicated number of HNRPNK eCLIP clusters (left) or the indicated number of SIRLOIN elements (right). **(F)** Changes in expression following HNRNPK knockdown in the indicated sample, comparing transcripts with either >2 or <2 eCLIP clusters. P-values computed using Wilcoxon test. **(G)** Changes in the indicated ratios for NucLibB fragments following HNRNPK knockdown. Each plot shows the difference between the ratio following HNRNPK siRNA transfection (average of two replicates) and non-targeting control siRNA transfection. **(H)** HNRNPK binding and the MLXIPL gene. Gene structure for a representative splicing isoform and SINE/LINE repeat annotations are from the UCSC genome browser. eCLIP reads/clusters are from HepG2 cells (replicate 1) from the ENCODE data portal. Expression levels in nucleus and cytoplasm in HepG2 and MCF7 cells from ENCODE and this study were normalized separately and HepG2 nuclear levels were capped to allow visual comparison. The exon enriched with HNRNPK binding is shaded.

We then tested whether HNRNPK binding was also associated with nuclear enrichment in transcripts that do not contain SIRLOIN elements. Strikingly, the number of HNRNPK binding clusters in eCLIP data correlated with stronger nuclear enrichment of lncRNAs and mRNAs in HepG2 (**Figure 4D**, Spearman R=0.14 P<10^-16^) and to a lesser extent in K562 cells (R=0.02, P=0.016, **Figure S9**), and this correlation was essentially identical when considering only transcripts that did not contain any SIRLOIN elements (R=0.153 for HepG2 cells). There was no correlation between nuclear enrichment and number of eCLIP binding peaks of a different poly(C) binding protein, PCBP2 (HepG2 cells, **Figure 4D**). This analysis suggests that HNRNPK takes part in a global pathway that mediates nuclear enrichment of long RNAs, and that is responsible for nuclear enrichment of transcripts with antisense Alu integrations.

To validate the role of HNRNPK in regulation of its bound targets, we knocked down HNRNPK in MCF7 cells using siRNAs (**Figure S10**), followed by RNA-seq of nuclear and cytoplasmic fractions as well as of whole cell lysate. In the control conditions, our fractionation data agreed very well with ENCODE data (R=0.76, P<10^-16^, **Figure S11**). Following HNRNPK knockdown, we observed a substantial effect on subcellular enrichment of hundreds of genes, with 397 genes becoming 2-fold more nuclear enriched and 283 genes becoming more cytoplasmic. Decrease in nuclear enrichment was significantly correlated with the number of HNRNPK eCLIP clusters (Spearman R=-0.22 between change in Nuc/Cyto ratio and number of eCLIP clusters, P<10^-16^, **Figure 4E**) and genes with >2 SIRLOIN elements were significantly less nuclearly enriched following HNRNPK knockdown (**Figure 4E**). Interestingly, when examining eCLIP-defined targets, changes in nuclear enrichment were mostly due to a reduction of transcript levels in the nucleus accompanied by only mild increase in cytoplasmic levels, overall resulting in slightly decreased expression levels of HNRNPK targets following knockdown (**Figure 4F**). We then tested the effect of HNRNPK knockdown on localization of transcripts from the NucLibB library. Consistently with the effects on endogenous targets, AcGFP with SIRLOIN-containing tiles became less enriched in the nucleus following HNRNPK depletion, predominantly through reduction in nuclear expression levels, with no significant changes in overall gene expression (**Figure 4G**). These results suggest that HNRNPK binding drives nuclear enrichment driven by SIRLOIN elements, but other factors likely contribute to decrease in overall expression levels associated with SIRLOIN integration.

The pathway uncovered here is likely relevant for at least some of the transcripts previously observed to be enriched in the nucleus *in vivo* in mice^3^. Overall, the nuclear/cytoplasmic ratios were strongly correlated between mouse liver and human HepG2 hepatocellular carcinoma cells (**Figure S13A**, Spearman R=0.51, P<10^-50^) and there was a significant correlation between number of HNRNPK eCLIP clusters in HepG2 and nuclear enrichment in the mouse liver (Spearman R=0.05, P=3.3·10^-7^). For instance, the transcription factor MLXIPL (also known as ChREBP) was previously shown to be retained in nuclear speckles in the mouse liver, beta cells, and intestine^3^. MLXIPL has a long internal exon containing multiple HNRNPK binding sites, is strongly enriched in the nucleus in various human cell lines, and was strongly affected by HNRNPK knockdown in MCF7 cells (**Figure 4H**).

Alu elements are primate-specific, but the mouse B1 repeat family is also derived from 7SL RNA and contains a sequence similar to the Alu SIRLOIN, and which we predict to also be regulated by HNRNPK. Indeed, Mlxipl#71 and Mlxipl#72 in NucLibA, two tiles from mouse Mlxipl 3' UTR that overlaps a B1 element in a negative-strand orientation, were associated with nuclear localization and reduced expression levels (**Figure S13B**). These tiles had the SIRLOIN-like sequence (TGG<u>CCTCC</u>AACTCAGAGATCCACCCA<u>CCCCT</u>G<u>CCTCT</u>GG) closer to the 3' end compared to other Mlxipl tiles, and only affected localization and expression when cloned into the 3' UTR, suggesting that the efficacy of this SIRLOIN-like B1-derived element, at least in human cells, is more restricted than that of human SIRLOIN. Both elements appear to have contributed to divergence in expression levels and localization between human and mouse – when homologous genes are compared (see Methods), transcripts that adopted SIRLOIN elements from Alus are more nuclear and lowly expressed in human hepatic cells, while those that adopted SIRLOIN-like B1 elements in mouse are more nuclear and lowly expressed in mouse (**Figure S13C**).

Since the HNRNPK consensus binding motif and the region we found to be essential for SIRLOIN activity in both JPX and PVT1 were C-rich motifs, we tested whether nuclear enrichment can be predicted from sequence alone. We counted the occurrences of all possible hexamers and correlated them with nuclear/cytoplasmic ratios in ENCODE cell lines. C-rich hexamers were among the best correlated with nuclear enrichment in human ENCODE cell lines (**Figure S14A-B**), and the same trend was observed also in mouse liver, mouse embryonic stem cells and MIN6 cells (using data from^3,18^), with the correlation typically stronger when considering occurrences only in internal exons (**Figure S14B**). This nuclear enrichment, predicted from sequence alone, was quite substantial – 311 genes expressed in HepG2 cells with at ≥50 occurrences of C-rich hexamers (>4 Cs) in their internal exons, and at least 2-fold more C-rich than G-rich motifs, were on average 3.4-fold more nuclear than genes with <50 occurrences (e.g, there are 84 C-rich hexamers in MLXIPL and only 10 of G-rich ones). These 311 genes with numerous C-rich hexamers were enriched for transcriptional regulators (GO categories “chromatin organization” and “positive regulation of transcription”, q-values 2.7·10^-4^ and 4.8·10^-3^, respectively, GOrilla analysis^19^), suggesting that such protein-coding genes, which are known to be generally less stable on both mRNA and protein levels^20^, are also more likely to be regulated by the HNRNPK-mediated nuclear enrichment pathway. Notably, a single consensus AluSx element in an antisense orientation has 44 hexamers with >3 Cs, and 5 with >4 Cs (compared to 14 with >3 Gs, and none with >4 Gs), and so integration of Alu elements in an antisense orientation substantially increases the C-rich hexamer content of the host transcript.

## Discussion

Alu elements are the most common SINE elements in the human genome, covering ~10% of the genome sequence^21^. Alus are enriched in transcribed regions and had a substantial impact on transcriptome evolution, for example through the contribution of new exons^22^ and polyadenylation sites^23^. Such events are actively suppressed in mRNAs^24,25^, but are common in lncRNAs^26,27^. Alu elements were also reported to act as functional modules in lncRNAs via intramolecular^7^ and intermolecular^28^ pairing with other Alus^29^. Here, we show that exonization of a specific part of an Alu element in an antisense orientation can affect the subcellular distribution of the RNA. SIRLOIN elements are quite common – 13.1% of lncRNAs and 7.5% of mRNAs in the human RefSeq database contain an Alu-derived SIRLOIN element, and 3.4% vs. 0.3% have a SIRLOIN element in an internal exon, which we find to be more effective. Exonization of Alus thus contributes to the tendency of lncRNAs to be enriched in the nucleus and expressed at lower levels when compared with mRNAs.

Our observation of repeated C/T-rich elements globally associated with nuclear enrichment and lower expression levels are also supported by previous studies of individual mRNAs and lncRNAs. For example, nuclear retention elements have been previously associated with decreased overall expression levels in β-globin mRNA^30^. An AGCCC motif has been reported to be important for nuclear retention of the BORG lncRNA^6^. A survey of known nuclear enrichment elements in ExportAid^31^ revealed several viral sequences containing closely spaced C/T hexamers, such as the Cis-acting Inhibitory Element (CIE) of the HTLV-1 and the two short polypyrimidine tracts associated with nuclear retention in HBV^32^. Viral RNAs may thus also rely on HNRNPK-mediated nuclear enrichment for maintaining low expression levels and nuclear retention during latency.

Multiplicity of sites and their context within the transcript appear to play a key role in determining the effectiveness of the HNRNPK binding sites. A large number of HNRNPK binding sites or C-rich hexamers correlates with stronger nuclear enrichment, and suggests additivity or even cooperativity between HNRNPK binding events in dictating localization. Other HNRNP proteins, such HNRNPA1, were shown to have strong cooperativity in RNA binding^33^. Several lines of evidence also suggest that the context of the HNRNPK binding sites influences their downstream effects. The position of SIRLOIN elements in NucLibB strongly correlated with activity, with elements closer to the 3' end of the transcript being more effective. C-rich hexamers and SIRLOIN elements were more strongly associated with nuclear enrichment when placed in internal exons.

Surprisingly, some well-studied nuclear lncRNAs, such as XIST and NEAT1 did not contain any regions which exhibited consistent nuclear-enrichment activity in our system, suggesting that their regions encoding nuclear enrichment are either longer than the 109 nt tiles, or are not active in our specific expression context. The Malat1 ~600 nt “region M” sequence^5^ causes strong nuclear enrichment in our system (**Figure 1E**), and another group found longer regions driving nuclear localization in XIST and NEAT1 (John Rinn, personal communication) and so we suggest that multiple independent pathways are likely responsible for nuclear enrichment in lncRNAs and mRNAs.

Assaying large numbers of synthesized or genome-derived sequences has been previously used for identifying elements driving transcription, splicing, RNA modifications, and other processes^34–36^. We report what is, to our knowledge, the first parallel screen for the identification of functional sequence elements in lncRNAs. The nuclear enrichment mechanism discovered in this screen is more common in lncRNAs, but also employed by some mRNAs, and so highlights the opportunities embodied in studying lncRNAs, which employ unique functional mechanisms and are under different selective pressures than mRNAs, to a general enhancement of our understanding of RNA biology. We expect that increasing understanding of the repertoire of lncRNA functions in cells will enable similar high-throughput approaches for identification of additional sequence and structural elements shared across lncRNAs, and expedite classification of these enigmatic genes into functional families.

## Methods

### Cell culture and transfection

MCF7 cells were grown in DMEM (Gibco) containing 10% fetal bovine serum and Penicillin-Streptomycin mixture (1%) at 37°C in a humidified incubator with 5% CO_2_.

Plasmids transfections were performed using PolyEthylene Imine (PEI)^37^ (PEI linear, M*r* 25000, Polyscience Inc). RNA was extracted 24hr following library transfections.

### Library design

Oligonucleotide pools were purchased from Twist Biosciences (San-Fransisco, CA). Tiles overlapping EcoRI, BglII, BamHI recognition sequences were excluded in both libraries and tiles overlapping HindIII, XbaI and NotI were also excluded in NucLibA.

### Plasmid library construction

Oligo pool was amplified by PCR (25ng template in 4.8ml reaction, divided into 96 50μl reactions, see **Table S10** for primer sequences), concentrated using Amicon tubes (UFC503096,millipore) and purified using AMpure beads (A63881, Beckman) at 2:1 beads:sample ratio according to manufacturer’s protocol. The insert was digested with EcoRI and BglII and cloned into likewise digested pAcGFP1-C1 or pAcGFP1-N1 (clontech) for 3’ and 5’ insertion respectively.

The ligation was transformed into E. coli electrocompetent bacteria (60117-2, lucigen) and plated on 15x15cm LB/Amp agar plates. Colonies were scraped off the plate and DNA was extracted using plasmid maxi kit (12163, Qiagen).

### Extraction of cytoplasmic and nuclear RNA

Cells were washed twice in cold PBS and resuspended in 300μl RLN buffer (50mM Tris•Cl pH8, 140mM NaCl, 1.5mM MgCl_2_, 10mM EDTA, 1mM DTT, 0.5% NP-40, 10U/ml RNAse inhibitor) and incubated on ice for 5 min. The extract was centrifuged for 5min at 300g in a cold centrifuge and the supernatant was transferred to a new tube, centrifuged again for 1min at 500g in a cold centrifuge, the supernatant (cytoplasmic fraction) was transferred again to a new tube and RNA was extracted using TRIREAGENT (MRC). The nuclear pellet was washed once in 300μl buffer RLN and resuspended in 1ml of buffer S1(250mM Sucrose, 10mM MgCl_2_, 10U/ml RNAse inhibitor), layered over 3ml of buffer S3 (880mM Sucrose, 0.5mM MgCl_2_, 10U/ml RNAse inhibitor), and centrifuged for 10min at 2800g in a cold centrifuge. The supernatant was removed and RNA was extracted from the nuclear pellet using TRIREAGENT.

### Sequencing library generation

One microgram of RNA was used for cDNA production using the qScript Flex cDNA synthesis kit (95049, Quanta) and a gene specific primer containing part of the Illumina RD2 region. The entire cDNA reaction was diluted into 100μ! second strand reaction with a mix of 6 primers introducing a unique molecular identifier (UMI) and a shift as well as part of the Illumina RD1 region. The second strand reaction was carried for a single cycle using Phusion HotStart polymerase (NEB), purified using AMpure beads at 1.5:1 beads: sample ratio, and eluted in 20μl of ddH_2_O. 15μl of the second strand reaction were used for amplification with barcoded primers, the amplified libraries were purified by twosided AMpure purification first with 0.6:1 beads to sample ratio followed by a 1:1 ratio.

NucLibA samples were sequenced with 119 nt reads and the NucLibB samples with 75 nt paired-end reads on an Illumina NextSeq 500 machine.

### Library data analysis

The sequenced reads were used to count individual library tiles using a custom Java script. We only considered R1 reads that contained the TTGATTCGATATCCGCATGCTAGC adapter sequence, and extracted the unique molecular identifier (UMI) sequence preceding the adapter. In R2 read, we removed the CGGCTTGCGGCCGCACTAGT adaptor and added the 3 bases preceding it to the UMI. Each read was then matched to the sequences in the library, without allowing indels. The matching allowed mismatches only at position with Illumina sequencing quality of at least 35 and we allowed up to two mismatches in the first 15 nt (“seed”), and no more than 4 overall mismatches. If a read matched more than one library sequence, the sequence with the fewest mismatches was selected, and if the read matched more than one library sequence with the same number of mismatches, it was discarded. Per-read mismatches were counted for the RNA editing analysis. See **Table S3** for read mapping statistics. The output from this step was a table of counts of reads mapping to each library sequence in each library (**Tables S3** and **S7**).

Only fragments with at least 10 reads on average in the WCE samples were used in subsequent analysis (5,153 fragments in NucLibA), and the number of UMIs mapping to each fragment was normalized to compute UPMs (UMI per million UMIs). We then used those to compute nuclear/cytoplasmic and WCE/input ratios after adding a pseudocount of 0.5 to each UPM (**Tables S4** and **S8**).

### Human transcriptome analysis

RefSeq transcript database (downloaded from the UCSC genome browser hg19 assembly on 30/06/2016) was used for all analyses. Only transcripts of exonic length of at least 200 nt were considered, and for entries mapping to multiple genomic loci, only one of the loci was used. We quantified the isoform-level expression levels using RSEM^38^ in the ENCODE data and computed the average fraction of each isoform among the isoforms of each gene. Only isoforms whose relative abundance was >25% were considered. Fold-changes between nuclear and cytoplasmic fractions were computed using DESeq2^39^. A transcript was considered to contain a SIRLOIN element or its antisense if it aligned with the sequence (without indels) with no more than eight mismatches.

### eCLIP data analysis

For alignment of eCLIP reads to Alu elements, We built a STAR aligner^40^ index using the JPX and Pvt1 Alu fragments, and aligned library reads to it using STAR with parameters: ‐‐outSAMstrandField intronMotif ‐‐readFilesCommand zcat ‐‐runThreadN 32 ‐‐outSAMtype BAM SortedByCoordinate ‐‐ outWigType bedGraph read1_5p. We then computed for each experiment the number of R2 reads whose 5' base mapped within the SIRLOIN fragment in either Pvt1 or Jpx sequence compared to the rest of the Alu sequence in each orientation.

In order to identify the sequence and structural preferences of the motif in HNRNPK eCLIP clusters, we used GraphProt^15^ to compare the eCLIP peaks to control peaks randomly sampled peaks of the same length from the same transcripts.

### RIP

RNA IP was performed according to the native RIP protocol described in Gagliardi & Matarazzo^41^. Anti-HNRNPK (RN019P, MLB) and Anti-GAPDH (C-2118S, Cell Signaling) were preincubated for 1h with protein-A/G magnetic beads (L00277, A2S) at 5μg antibody/ 50μl bead slurry. Extract from 2x10^6^ cells was added to the beads and incubated overnight rotating at 4°c. Precipitated RNA was extracted and analyzed by RT-qPCR (see **Table S10** for primers).

### Imaging Flow Cytometry

MALAT1 and AcGFP mRNAs were labeled using the PrimeFlow RNA Assay kit (Affymetrix) according to the manufacturer’s protocol and visualized using the ImageStream®^X^ Mark II (Amnis).

### RNA-seq

MCF7 cells were transfected with 10nM siRNA pool targeting HNRNPK or with control pool (Dharmacon) using Lipofectamin 3000 reagent (L3000001, thermo-fisher), RNA was extracted 72 hrs post-transfection and libraries were prepared using the SENSE mRNA-Seq Library prep kit (SKU: 001.24, Lexogen) according to manufacturer’s protocol.

RNA-seq reads were mapped to the mouse genome (mm9 assembly) with STAR^40^ and transcript levels were quantified using RSEM^38^ with default parameters. Fold-changes were computed using DESeq2^39^.

### HNRNPK knockdown in cells expressing NucLibB

MCF7 cells were transfected with siRNA pool or non-targeting control as described above. 48 hours post transfection the cells were washed and transfected with NucLibB plasmids. RNA was extracted after an additional 24 hours and libraries were prepared as described above. Libraries were analyzed the same way as described above and nuclear/cytoplasmic, nuclear/input, cytoplasmic/int and WCE/input ratios after adding a pseudocount of 0.5 to each UPM. Ratios for the two HNRNPK siRNA replicates were averaged.

### Conservation analysis

RefSeq transcripts for which we quantified nuclear/cytoplasmic ratios were mapped to Ensembl transcripts and human and mouse orthologs were obtained from Ensembl Compara 80. Nuclear/cytoplasmic ratios were compared between HepG2 (ENCODE data) and mouse liver^3^ and expression levels between human liver expression (HPA data^42^) and ENCODE mouse liver expression, both quantified using RSEM. Only genes with FPKM≥0.5 in at least one of the datasets were considered for the analysis. Overlaps with Alu and B1 elements were computed using RepeatMasker data from the UCSC genome browser.

### Hexamer analysis

Sequences for RefSeq transcripts described above were extracted from the UCSC genome browser. Occurrences of all possible hexamers (allowing overlaps) were counted in either all the exons, or just in the internal or terminal exons. Then, for each hexamer, the Spearman correlation coefficient was computed between the total number of occurrences of the motif and the nuclear/cytoplasmic ratio as computed by DESeq2.

## Accession numbers

All sequencing data has been deposited to the SRA database, accession SRPXXXX.

## Author contributions

YL and IU conceived and designed the study. YL carried out all experiments and IU carried out computational analysis. YL and IU wrote the manuscript.

